# The association between the *HLA-DRB1* shared epitope alleles and the risk of rheumatoid arthritis is influenced by massive gene-gene interactions

**DOI:** 10.1101/221416

**Authors:** Lina-Marcela Diaz-Gallo, Daniel Ramsköld, Klementy Shchetynsky, Lasse Folkersen, Karine Chemin, Boel Brynedal, Steffen Uebe, Yukinori Okada, Lars Alfredsson, Lars Klareskog, Leonid Padyukov

## Abstract

In anti-citrullinated protein antibody positive rheumatoid arthritis (ACPA-positive RA), a particular subset of *HLA-DRB1* alleles, called shared epitope alleles (SE), is the highest genetic risk factor. Here, we aimed to investigate whether gene-gene interactions influence this *HLA-DRB1* related major disease risk; specifically, we set out to test if non-HLA SNPs, conferring low diseases risk on their own, can modulate the *HLA-DRB1* SE effect to develop ACPA-positive RA.

To address this question, we computed the attributable proportion (AP) due to additive interaction at genome-wide level for two independent ACPA-positive RA cohorts: the Swedish EIRA and the North American NARAC. We found a strong enrichment of significant interactions (AP p-values<0.05) between the *HLA-DRB1* SE alleles and a group of SNPs associated with ACPA-positive RA in both cohorts (Kolmogorov-Smirnov [KS] test D=0.35 for EIRA and D=0.25 for NARAC, p<2.2e-16 for both). Interestingly, 201 out of 1,492 SNPs in consistent interaction for both cohorts, were eQTLs in SE alleles context in PBMCs from ACPA-positive RA patients. Finally, we observed that the effect size of *HLA-DRB1* SE alleles for disease decreases from 5.2 to 2.5 after discounting the risk alleles of the two top interacting SNPs (rs2476601 and rs10739581, AP FDR corrected p <0.05).

Our data demonstrate that the association between the *HLA-DRB1* SE alleles and the risk of ACPA-positive RA is modulated by massive genetic interactions with non-HLA genetic variants.

## Introduction

Additive interaction, defined as the deviation from the expected sum of the effects of two different factors, is a way to explore the complexity of how individual genetic risk variants interplay in the development of complex diseases. However, the possibility to address these additive interactions between candidate variants is often limited by low statistical power. Additionally, genome-wide gene-gene interaction studies conceivably result in a high number of false negative results due to the massive and conservative correction for multiple testing. An alternative strategy to study interaction is to identify genetic “hubs” that may accumulate multiple interactions with different variants. As a result of these interactions, such genetic “hubs” may have a strong influence on the risk of disease.

In rheumatoid arthritis (RA [OMIM: 180300]), a particular subset of *HLA-DRB1* gene variants (major alleles at *01, *04, and *10 groups), commonly called shared epitope (SE) alleles, is the most important genetic contributor for the risk of developing anti-citrullinated protein antibody (ACPA) positive RA (1, 2). It is noteworthy that the strength of the association between non-*HLA* genetic variants and ACPA-positive RA risk is, in general, very moderate in comparison to that of the *HLA-DBR1* SE alleles (3–6), (Fig. 1a). This prompted us to suggest that the *HLA-DRB1* SE alleles could be a genetic “hub” that captures multiple interactions. Indeed, previous studies have demonstrated interactions between the *HLA-DRB1* SE alleles and several SNPs, including variations in *PTPN22, HTR2A*, and *MAP2K4* with regard to the risk of developing ACPA-positive RA (7–10), where the combination of both risk factors shows significantly higher risk (measured as odds ratio (OR)) than the sum of their separate effects. Departure from additivity is a way to define and subsequently demonstrate interaction between risk factors regarding the risk of disease. The additive scale, defined by attributable proportion (AP), has the advantage of a straightforward interpretation in the sufficient-component cause model framework (7, 11–14).

**Fig 1.**
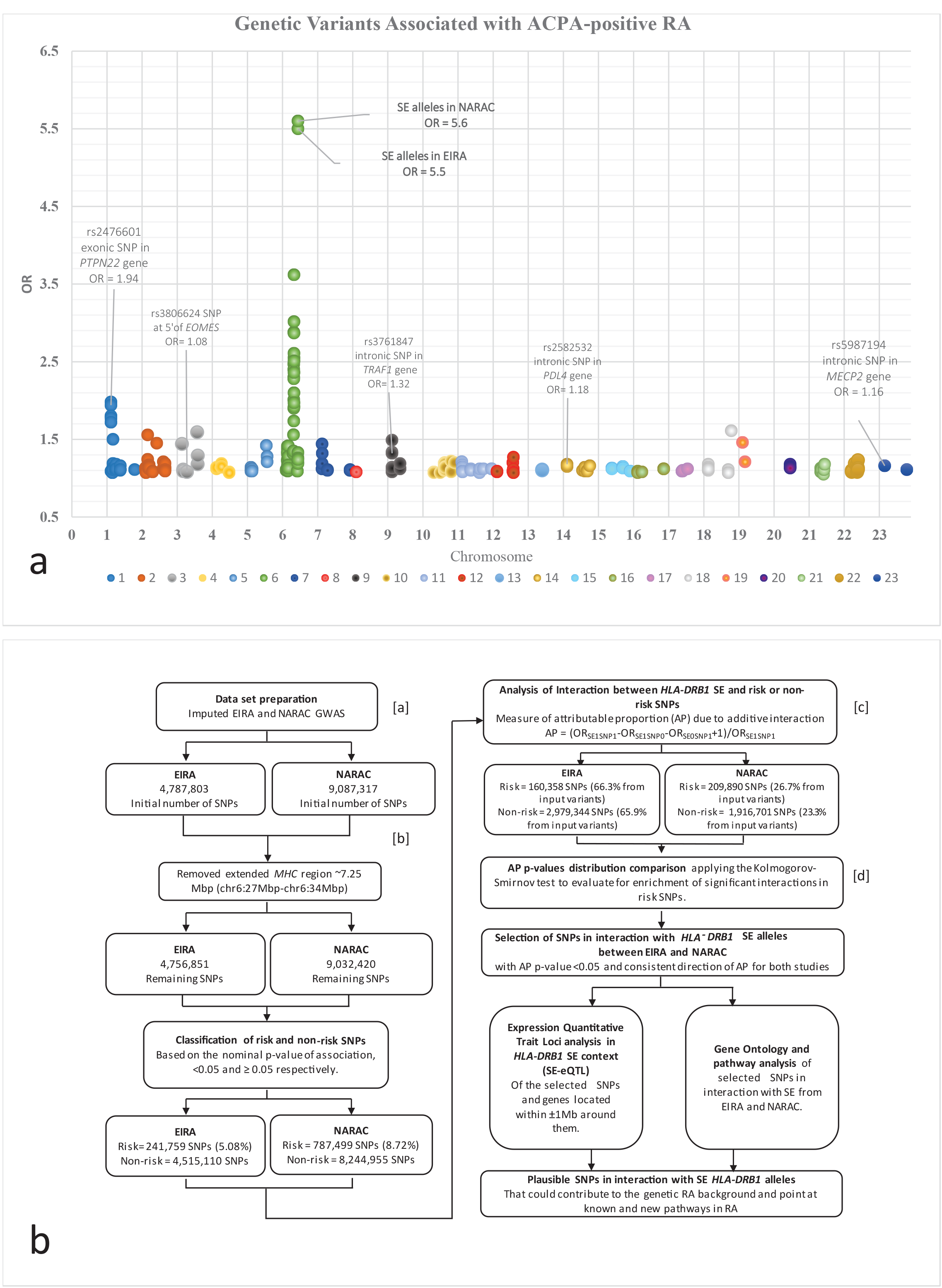
(a) Genetic variants associated with ACPA-positive RA. This plot represents the association signals (p-values < 1.0e-05) from different GWAS in ACPA-positive RA, taken from the NHGRI-EBI GWAS catalog (<u>https://www.ebi.ac.uk/gwas/home</u>) (50–52). The x-axis shows the physical positions for the chromosomes of the human genome including chromosome X (marked as 23). The y-axis represents the OR value observed for each SNP in different studies. As examples, some polymorphisms are pointed out together with the OR observed for the *HLA-DRB1* SE alleles in EIRA and NARAC studies. **(b) Methodology work flow**. The workflow applied in the present study. **[a]** The same workflow was applied using only genotyping data (non-imputed) from both cohorts. In an independent analysis, all genetic variations from the *MHC* locus or from the entire chromosome 6 were removed from the analysis (Supplementary Material Table S2). **[b]** An alternative step was included at this point of the workflow. The *PTPN22* locus (chr1:113679091 to chr1:114679090) was removed from the analysis due to the previously reported interaction in RA between the non-synonymous variant rs2476601 in the *PTPN22* gene and the *HLA-DRB1* SE alleles (7). **[c]** The gender and the first ten principal components from PCA were included as covariates in the model. The interaction test was only applied when at least 5 individuals were present in each combined category of the calculation. AP value, its respective p-value and confidence interval (95%CI) were assessed using logistic regression by means of the program GEISA (https://github.com/menzzana/geisa)(11, 40). **[d]** In order to verify that the observed results were not due to statistical artifacts, we employed several approaches. First, we permuted the category of risk and non-risk for the observed AP p-values ten thousand times, applying the KS test each time to identify the proportion of p-values from the KS tests that are less than the set threshold (<2.2e-16). The percentage of the KS test results with p-values less than 2.2e-16 was 0, the maximum D value observed was 6.1e-03 in both analyzed cohorts. Likewise, we permuted the *HLA-DRB1* SE variable using non-imputed GWAS data, with a smaller number of permutations (n=1000) due to the computational limitations, to calculate the interaction for each randomized SE variable against all SNPs in the GWAS. We applied the KS test to detect differences in the AP p-values’ distribution for risk (nominal p-values of association <0.05) versus non-risk (nominal p-values of association >0.05) SNPs, each time the SE variable was randomized. The maximum D values observed were 0.05 and 0.03 for EIRA and NARAC, respectively. The percentage of the KS test p-values from permutations less than 2.2e-16 were 0.1 for both EIRA and NARAC. Both types of permutations showed that less than 5% of the KS test will exhibit a p-value under 2.2e-16, strongly indicating that differences in the AP p-values distribution detected by the KS test from the original data are unlikely to be by chance. Secondly, as it is mentioned before, we tested the same workflow after removing the *PTPN22* locus, to test whether the enrichment of interactions observed was significantly influenced by these variants. Third, we applied the same workflow up to this point, replacing the SE variable with the rs4507692 SNP as a negative control, since the rs4507692 SNP is not associated with ACPA-positive RA but has the same MAF as the *HLA-DRB1* SE alleles. Fourth, as it is mentioned before, non-imputed GWAS data were used in the same methodological workflow, as well as removing data from the entire chromosome 6, to address possible inflation in the results due to a high LD with the *HLA-DRB1* SE alleles. Abbreviations: SE1SNP1: presence of the *HLA-DRB1* SE alleles and the risk allele from the SNP, SE1SNP0: presence of the *HLA-DRB1* SE alleles and absence of the risk allele from the SNP, SE0SNP: absence of the *HLA-DRB1* SE alleles and presence of the risk allele from the SNP, ACPA-positive RA – anti-citrullinated protein antibody positive rheumatoid arthritis, SE: share epitope, GWAS – genome-wide association study, NHGRI – National Human Research Institute, EBI – European Bioinformatics Institute, OR – odds ratio, EIRA – epidemiological investigation of rheumatoid arthritis, NARAC – North American rheumatoid arthritis consortium, *MHC* locus – major histocompatibility locus, PTPN22 – gene abbreviation, PCA – principal component analysis, KS – Kolmogorov-Smirnov test, MAF – minor allele frequency, LD – linkage disequilibrium.

In our current study, we aimed to investigate whether gene-gene interactions influence the major *HLA-DRB1* related disease risk to develop ACPA-positive RA; more specifically, we set out to test if non-HLA SNPs, conferring low diseases risk on their own, can modulate the *HLA-DRB1* SE effect to develop ACPA-positive RA. First, we assessed departure from additivity regarding the interaction between the *HLA-DRB1* SE alleles and SNPs at the genome-wide level. The outcome of this analysis was tested for the enrichment of significant interactions by comparing the distribution of studied statistics (p-value of interaction) between two defined groups of SNPs: the pool of SNPs which exhibited a significant nominal association with ACPA-positive RA in comparison to SNPs that are not associated with disease risk. Second, we performed the same type of analysis in an independent ACPA-positive RA cohort in order to replicate our findings. Third, we analyzed the effect size from the *HLA-DRB1* SE alleles with regard to risk of ACPA-positive RA before and after step-by-step discount of the risk alleles of the strongest SNPs in interaction with SE. Finally, we performed an expression quantitative trait loci (eQTL) analysis stratifying by the *HLA-DRB1* SE alleles and pathway enrichment analysis, in a further step to contextualize the selected SNPs in interaction with the *HLA-DRB1* SE alleles from both studied cohorts. Our observations indicated that the effect of the *HLA-DRB1* SE alleles in the development of ACPA-positive RA is influenced by interactions with multiple non-HLA genetic factors, supporting the concept that these *HLA-DRB1* alleles act as a “hub” of cumulative additive interactions with multiple genetic variants. We proposed with the present methodology a novel approach to study the impact of gene-gene interactions with *HLA* alleles in autoimmune diseases.

## Results

This project was based on genome wide association studies (GWAS) data from two independent case control studies of RA, Epidemiological Investigation of Rheumatoid Arthritis (EIRA) (5, 12, 15–18) and North American Rheumatoid Arthritis consortium (NARAC) (5, 16, 19, 20). The overall methodology workflow is shown in Fig. 1b. We assessed pair additive interactions, measured by AP, between the *HLA-DRB1* SE alleles and *non-HLA* SNPs in EIRA and NARAC. We used the described workflow (Fig. 1b) in parallel to both the original genotyped data sets, the imputed data sets and the rs4507692 SNP instead of the *HLA-DRB1* SE alleles. The rs4507692 was considered as a negative control, since this variant exhibit the same minor allele frequency (MAF) as *HLA-DRB1* SE alleles but is not associated to ACPA-positive RA (Table 1). We tested for enrichment of significant interactions between two predefined groups of SNPs, the ACPA-positive RA risk SNPs (nominal p-value of association < 0.05) and the ACPA-positive RA non-risk SNPs (nominal p-value of association ≥ 0.05) using the Kolmogorov-Smirnov (KS) test. The KS test statistic quantifies the maximum distance (D) between the two empirical cumulative distribution functions (ECDF) of the AP p-values from the risk and nonrisk SNPs groups.

**Table 1.**
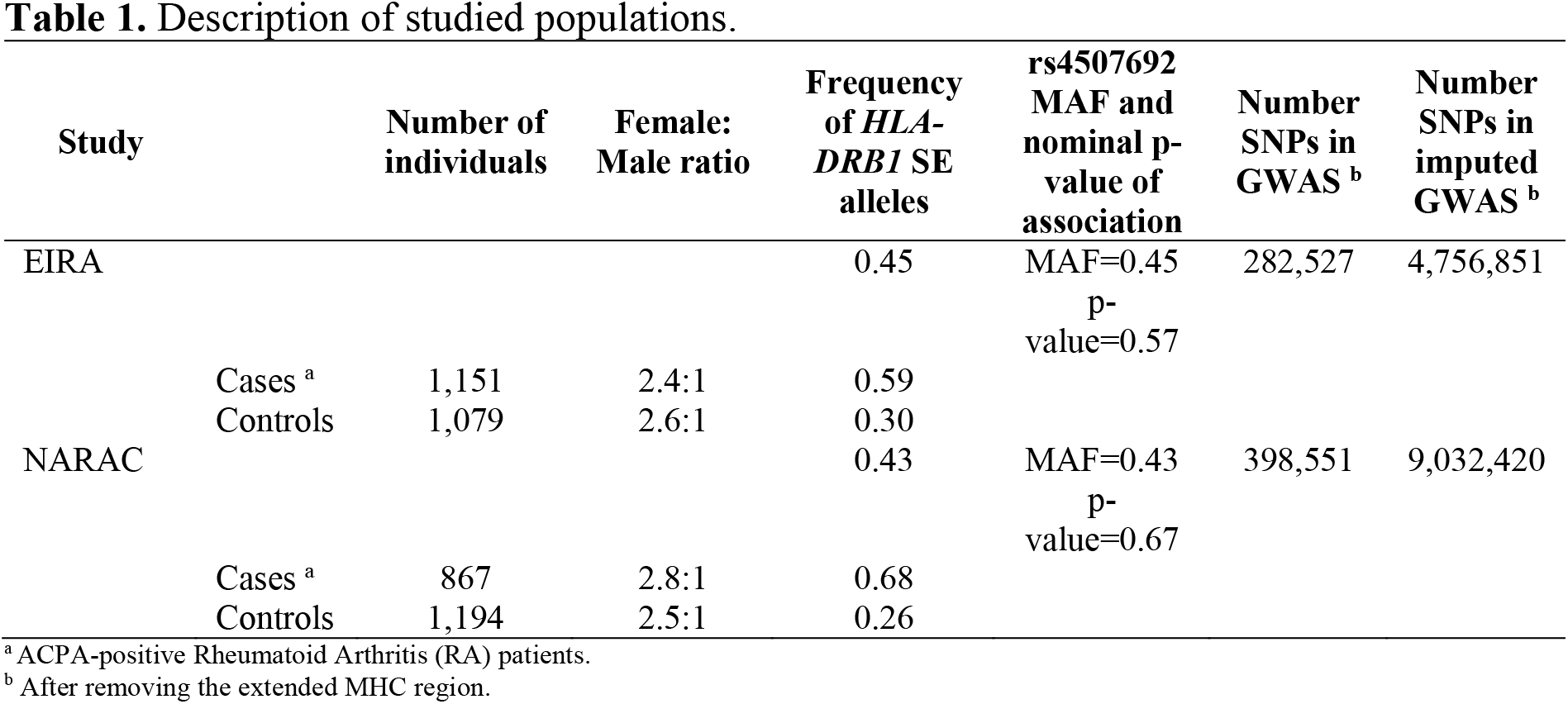
Description of studied populations.

### Interaction of the HLA-DRB1 SE alleles with ACPA-positive RA associated SNPs is more common than with non-associated SNPs

EIRA was considered as a discovery cohort to test for enrichment of significant interactions between the *HLA-DRB1* SE alleles and the set of SNPs enriched for risk SNPs from this study. The risk SNPs represent 5% of the variants analyzed for interaction in EIRA. Out of these risk SNPs, 24.5% of them exhibited an AP p-value (attributable proportion due to interaction p-value) less than 0.05 (Table 2, Fig. 2a). On the other hand, among the non-risk variants (nominal p-values of association ≥0.05) representing the remaining SNPs analyzed for interaction in EIRA, only 2.8% displayed a significant interaction (AP p-value <0.05) with the *HLA-DRB1* SE alleles (Table 2, Fig. 2b). Thus, there is a dramatic difference in the frequency of significant interactions with the *HLA-DRB1* SE alleles between the risk and non-risk SNPs in ACPA-positive RA. This observation is reflected in the KS test, where a striking difference was observed between the AP p-values’ distributions of risk and non-risk SNPs with a D value of 0.35 (KS test p-value <2.2e-16) (Table 2 and Fig. 2c).

**Fig 2.**
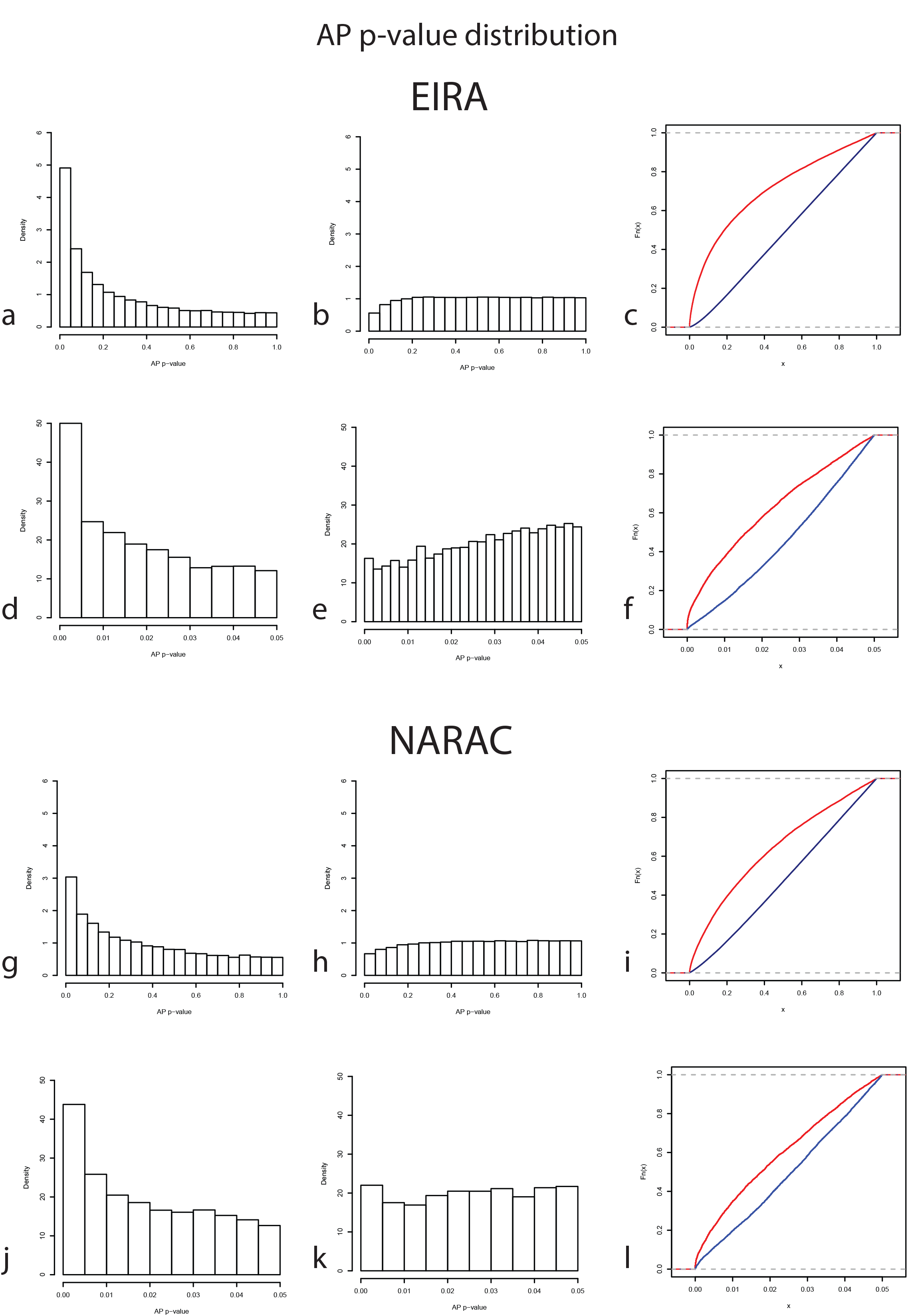
Comparison of the distribution of p-values for attributable proportion in EIRA and NARAC studies for interaction tests between the *HLA-DRB1* SE alleles and genetic variants. (a) Density plot of AP p-values for the interaction between the *HLA-DRB1* SE alleles and the risk group of SNPs (nominal p-value of association <0.05) or **(b)** non-risk group of SNPs (nominal p-value of association ≥0.05) in the EIRA study. **(c)** The respective ECDF plot of the AP p-values distribution of risk (red line) or non-risk (blue line) SNPs in interaction with the *HLA-DRB1* SE alleles (KS test, D=0.35, p-value <2.2e-16; Table 2). We tested for differences in the AP p-values distribution on the fraction that could be considered as significant interactions with the *HLA-DRB1* SE alleles (AP p-value <0.05). **(d)** Density plot for the AP p-values from the interaction tests between the risk SNPs and the *HLA-DRB1* SE alleles or, **(e)** between the non-risk SNPs and the *HLA-DRB1* SE alleles in the EIRA study. **(f)** ECDF of the fraction of AP p-values distribution corresponding to <0.05 in the EIRA study (KS test, test D=0.26, p-value >2.2e-16). Similar results were observed from the NARAC study, an independent replication cohort: **(g)** Density plot of the AP p-values for the interaction between the *HLA-DRB1* SE alleles and the risk group of SNPs or **(h)** non-risk group of SNPs. **(i)** The respective, ECDF plot from the NARAC study (KS test, D=0.25, p-value <2.2e-16, Table 1). **(j)** Density plot of the fraction of the AP p-values distribution of less than 0.05 from the interactions between the *HLA-DRB1* SE alleles and the risk SNPs or **(k)** non-risk SNPs. **(l)** The ECDF plot from this fraction of the AP p-values distribution (KS test, D=0.17, p-value >2.2e-16). Abbreviations: EIRA – epidemiological investigation of rheumatoid arthritis, NARAC – North American rheumatoid arthritis consortium, AP – attributable proportion due to interaction, ECDF – Empirical cumulative distribution function, KS test – Kolmogorov – Smirnov test.

**Table 2.** The Kolmogorov-Smirnov (KS) test for AP p-values distributions of the interaction analysis with the *HLA-DRB1* SE alleles and risk or non-risk SNPs in EIRA and NARAC imputed data.

| Case-Control Group | SNPs group | Number of initial input SNPs | Number of SNPs after cut off <sup>a</sup> | % of SNPs analyzed | Number of SNPs with AP p-value $< 0.05$ | % of analyzed SNPs with AP p-value $< 0.05$ | D<sup>+</sup> value from KS test<sup>b</sup> | Group of SNPs with enrichment of significant interactions |
| --- | --- | --- | --- | --- | --- | --- | --- | --- |
| EIRA | Risk | 241,759 | 160,358 | 66.33 | 39,518 | 24.64 | 0.354 | Risk |
|  | No-risk | 4,515,110 | 2,979,344 | 65.99 | 83,287 | 2.80 |  |  |
| NARAC | Risk | 787,499 | 209,890 | 26.65 | 31,992 | 15.24 | 0.247 | Risk |
|  | No-Risk | 8,244,955 | 1,916,701 | 23.25 | 64,012 | 3.44 |  |  |

<sup>a</sup> Interaction was done using sex and the 10 first eigenvectors as co-variables. Cutoff used a minimum of 5 individuals for each of the OR combinations.
<sup>b</sup> The alternative hypothesis for KS test (Kolmogorov-Smirnov test) was that the ECDF (empirical cumulative distribution function) of AP p-values for risk SNPs lies above that of non-risk SNPs (Figure 2). KS test p-value $< 2.2 \times 10^{-16}$ for both EIRA and NARAC. As it is mentioned in the materials and methods, these KS test p-values are lower than the machine precision, meaning that when the precise p-value was calculated the result was 0.

The difference in the distribution of the whole spectrum of AP p-values does not directly inform about the performance of the values below significance threshold of 0.05. Therefore, we specifically tested for difference in this segment of the AP p-values distributions for risk and non-risk SNPs. We found a strong enrichment of significant interactions in the group of risk variants in comparison with the non-risk variants (KS test D=0.25, p-values <2.2e-16, Fig. 2d to Fig. 2f). This suggests that the significant difference between the ECDF from the risk and non-risk groups detected in the full distribution of AP p-values is principally due to the enrichment of small AP p-values in the risk group of SNPs.

Since genetic variants located in the *PTPN22* gene are the second most important genetic risk factor for RA in Caucasians (Fig. 1a), we excluded the SNPs of this locus from the analysis and tested for the enrichment of significant interactions between the *HLA-DRB1* SE alleles and the risk group of SNPs. The exclusion of the *PTPN22* locus did not remarkably affected the obtained D and p-values from the KS test (D=0.353, p-value<2.2e-16). This highlights that the enrichment of significant interactions between the ACPA-positive RA risk SNPs and the *HLA-DRB1 SE* alleles is due to multiple variants, and it is not explained by the *PTPN22* locus alone.

The ECDF difference between the AP p-values from the risk and non-risk variants almost disappeared completely when the rs4507692 SNP was tested instead of the *HLA-DRB1* SE variable as a negative control (Table 1, Supplementary Material Table S1). We found that the proportion of interacting risk SNPs with rs4507692 variant dropped to 2.8%, (Supplementary Material Table S1 and Fig. S1a to Fig. S1f). Since the same group of risk variants was tested for interaction with the *HLA-DRB1* SE alleles and rs4507692 SNP, we evaluated for differences in the AP p-value distributions between both set of analyses. This analysis confirmed that there is a high enrichment of significant interactions between the risk variants and the *HLA-DRB1*

SE alleles (D value=0.35, KS-test p-value< 2.2e-16, Supplementary Material Fig. S2a). These results demonstrate that the enrichment of interactions found for the *HLA-DRB1* SE alleles is unlikely due to a random effect and point to the specific role of the *HLA-DRB1* SE alleles in the architecture of gene-gene interaction in ACPA-positive RA.

Consistent results were observed when the workflow was applied to only non-imputed genotyping data for EIRA (Supplementary Material Table S2). Additionally, we also removed all chromosome 6 markers (where the *MHC* region lies) to exclude any influence of LD with the *HLA-DRB1* SE alleles in this chromosome. In this analysis, we observed an enrichment of significant interactions between the *HLA-DRB1* SE alleles and the risk variants, either when only the *MHC* region was removed (KS-test D=0.33, p-value<2.2e-16, Supplementary Material Table S2) or when the entire chromosome 6 was removed (KS-test D=0.33, p-value<2.2e-16, Supplementary Material Table S2). No significant differences in the ECDF of AP p-values were observed when the rs4507692 SNP was implemented in the workflow instead of the *HLA-DRB1* SE alleles (KS-test D=0.006, p-value=0.39 for non-imputed EIRA GWAS without the MHC region and KS-test D=0.007, p-value=0.32 for the non-imputed SNPs GWAS without the chromosome 6, Supplementary Material Table S2). These results indicate that high LD variants present in the imputed GWAS data set do not inflate the difference between the ECDF of AP p-values from the interaction analysis of the *HLA-DRB1* SE alleles and groups of risk or nonrisk SNPs.

### An independent replication supports the observed enrichment of significant interactions between the HLA-DRB1 SE alleles and the ACPA-positive RA associated SNPs

In order to confirm the results observed in the EIRA study, we applied the same methodology in the independent case-control NARAC study. Similar to EIRA, we found a higher enrichment of significant interactions between the *HLA-DRB1* SE alleles and the risk SNPs (15.2%) in comparison to the significant interactions detected between the *HLA-DRB1* SE alleles and the non-risk SNPs (3.3%) (Table 2, Fig. 2g and Fig. 2h). The KS test reflected such a difference in the ECDF of the AP p-values, with a D value of 0.25 (p-value <2.2e-16, Table 2, Fig. 2i). Similar to our findings in the discovery cohort, the fraction of AP p-values below 0.05 is enriched in the risk group of SNPs compared to the non-risk group of variants in the NARAC study (D=0.17, p-value <2.2e-16, Fig. 2j to Fig. 2l). As in EIRA, when the rs4507694 SNP was used in the workflow instead of the *HLA-DRB1* SE alleles, there was not an enrichment of significant interactions in the risk group (2.6%) compared to the non-risk group (3%) of SNPs (Supplementary Material Table S1, Fig. S1g to Fig. S1l). Also, the distribution of AP p-values from the *HLA-DRB1* SE alleles and the risk SNPs is strongly different from the distribution of the AP p-values from the rs4507692 (with the same MAF as SE, but not associated to ACPA-positive RA) and the risk SNPs (KS test D=0.26, p-value<2.2e-16; Supplementary Material Fig. S2b). Consistent results were observed when genotyped sets of SNPs (non-imputed GWAS) were used for the analyses in the NARAC study (Supplementary Material Table S2).

### Step-by-step discount of the risk alleles of top interacting SNPs decreases the HLA-DRB1 SE risk for ACPA-positive RA

The definition of additive interaction model predicts that removing individuals with an interacting allele from the analysis should decrease the effect size of the *HLA-DRB1* SE alleles among the remaining subjects. To directly determine if the OR for the *HLA-DRB1* SE alleles in ACPA-positive RA is affected by the absence or presence of other risk alleles from SNPs in interaction with the *HLA-DRB1* SE alleles, we calculated the combined OR for the *HLA-DRB1* SE alleles, including and excluding the effect of two of the top SNPs in interaction. Figure 3 shows how the combined OR is affected by the exclusion of individuals with one or both of the risk alleles of the selected SNPs in combination with the *HLA-DRB1* SE alleles. Importantly, in EIRA, removing the risk alleles gradually decreases the ORs from 11.48 (8.9–14.9 95%CI; YGG: SE positive, rs2077507G, rs1004664G) to 3.97 after removing one risk allele (95%CI: 3.3 – 4.7; YAG: SE positive, rs2077507A, rs1004664G) or to 4.11 after removing the other risk allele (95%CI: 3.3 – 5.1; YGT: SE positive, rs2077507G, rs1004664T) and finally, to an OR of 2.57 when both risk alleles are removed (95%CI: 2.2 – 2.9; YAT: SE positive, rs2077507A, rs1004664T, Fig. 3a). A similar result was seen in the NARAC study for the top interacting SNPs (Fig. 3b). This gradual drop in effect size indicates that the high OR of the *HLA-DRB1* SE alleles in the disease could be at least partially attributed to the interaction with other SNPs, which exhibit modest individual effect on the risk for ACPA-positive RA.

**Fig 3.**
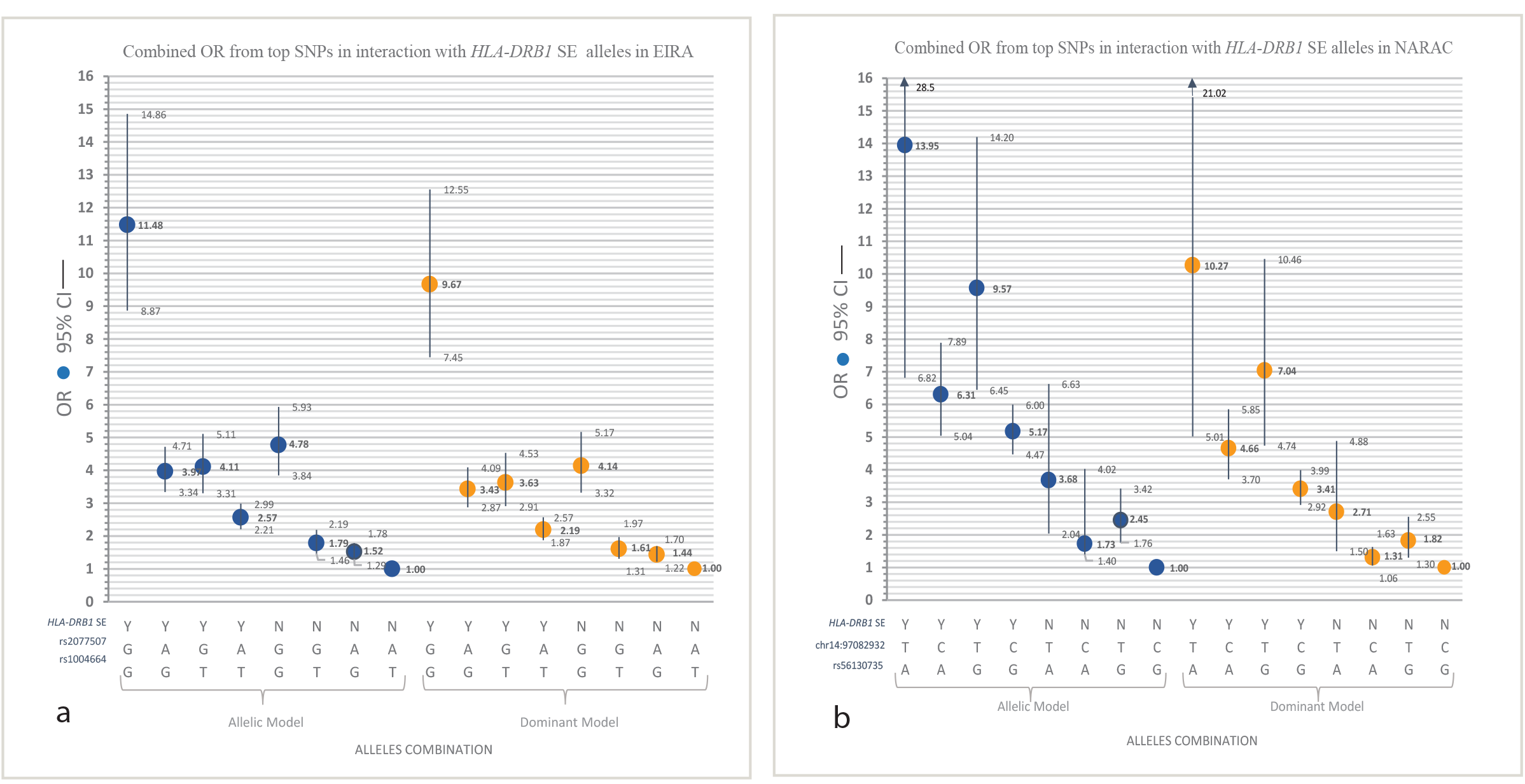
Three-factor’s OR calculation: the *HLA-DRB1* SE alleles and the two most significant SNPs in interaction with each cohort. On the x-axis – the combinations of presence or absence of risk alleles for allelic or dominant models. On the y-axis – the combined ORs with 95% CI of *HLA-DRB1* SE alleles (presence – Y, or absence – N) and the most significant SNPs in interaction from each cohort. Panel **(a)** shows data from the EIRA study, where the rs2077507(A>G) and rs1004664(T>G) SNPs are represented. The YGG alleles combination represents all risk alleles. The rs2077507 SNP is in significant interaction with the *HLA-DRB1* SE alleles (AP=0.57 95%CI=0.47–0.67, p-value <1e-16). Similarly, rs1004664 is in significant interaction with the *HLA-DRB1* SE alleles (AP=0.46 95%CI=0.34–0.57, p-value=6e-14) in the EIRA study. Panel **(b)** shows data from the NARAC study, where the chr14:97082932(C>T) and rs56130735(G>A) variants are represented. The YTA alleles combination represents all risk alleles. The chr14:97082932 variant shows significant interaction with the *HLA-DRB1* SE alleles (AP=0.54 95%CI=0.34–0.75, p-value=2.1e-07) as well as rs56130735 (AP=0.37 95%CI=0.19–0.54, p-value=4.7e-05) in the NARAC study. Abbreviations: OR – odds ratio, CI – confidence intervals, EIRA – epidemiological investigation of rheumatoid arthritis, NARAC – North American rheumatoid arthritis consortium, AP – attributable proportion due to interaction.

### An exploration of selected SNPs in interaction with the HLA-DRB1 SE alleles from EIRA and NARAC

We identified 1,492 SNPs in interaction with the *HLA-DRB1* SE alleles with AP p-values <0.05 and the same direction of AP when comparing the results from the EIRA and NARAC studies (Supplementary Material Table S3).

Figures 4a and 4b visualize how these 1,492 SNPs are distributed across the genome. We ranked the chromosomes based on the minimum AP p-value, the maximum AP value, and the percentage of these 1,492 SNPs in interaction with the *HLA-DRB1* SE alleles (Supplementary Material Table S4). Based on these criteria, chromosomes 1 and 9 reach the highest position for both studied cohorts (minimum AP p-value 4.3e-10 in EIRA and 1.6e-08 in NARAC; Supplementary Material Table S4). Chromosomes 2, 7, 8, and 13 followed in the ranking when the results from both EIRA and NARAC were considered. The majority (84.6%) of these SNPs in interaction with the *HLA-DRB1* SE alleles exhibited a positive AP, and most of them had values under 0.5 (Fig. 4a and Fig. 4b). The genotypes of 201 variants out of 1,492 (13.5%) were statistically significant correlated with the expression of different genes located 2Mb around them, in peripheral blood mononuclear cells (PBMCs) from the ACPA-positive RA patients, when the *HLA-DRB1* SE alleles stratification was applied (SE-eQTLs). Supplementary Material Table S5 contains a complete list of the SNP-gene pairs that exhibited a false discovery rate (FDR) q-value< 0.05 for the SE-eQTLs analysis. Among the top SE-eQTLs are rs10404242-TLE6 (transducing like enhancer of split 6) at chromosome 19, rs5763638-ZNRF3-*AS1* (ZNRF3 antisense RNA 1) at chromosome 22, rs28513183-HSD11B1 (hydroxysteroid 11-beta dehydrogenase 1) at chromosome 1, and rs1781279-MTPAP (mitochondrial poly(A) polymerase) at chromosome 10 (Supplementary Material Fig S3). Since these SE-eQTLs are context related, it gives biological evidence for the statistically detected interactions between these variants and the *HLA-DRB1* SE alleles.

**Fig 4.**
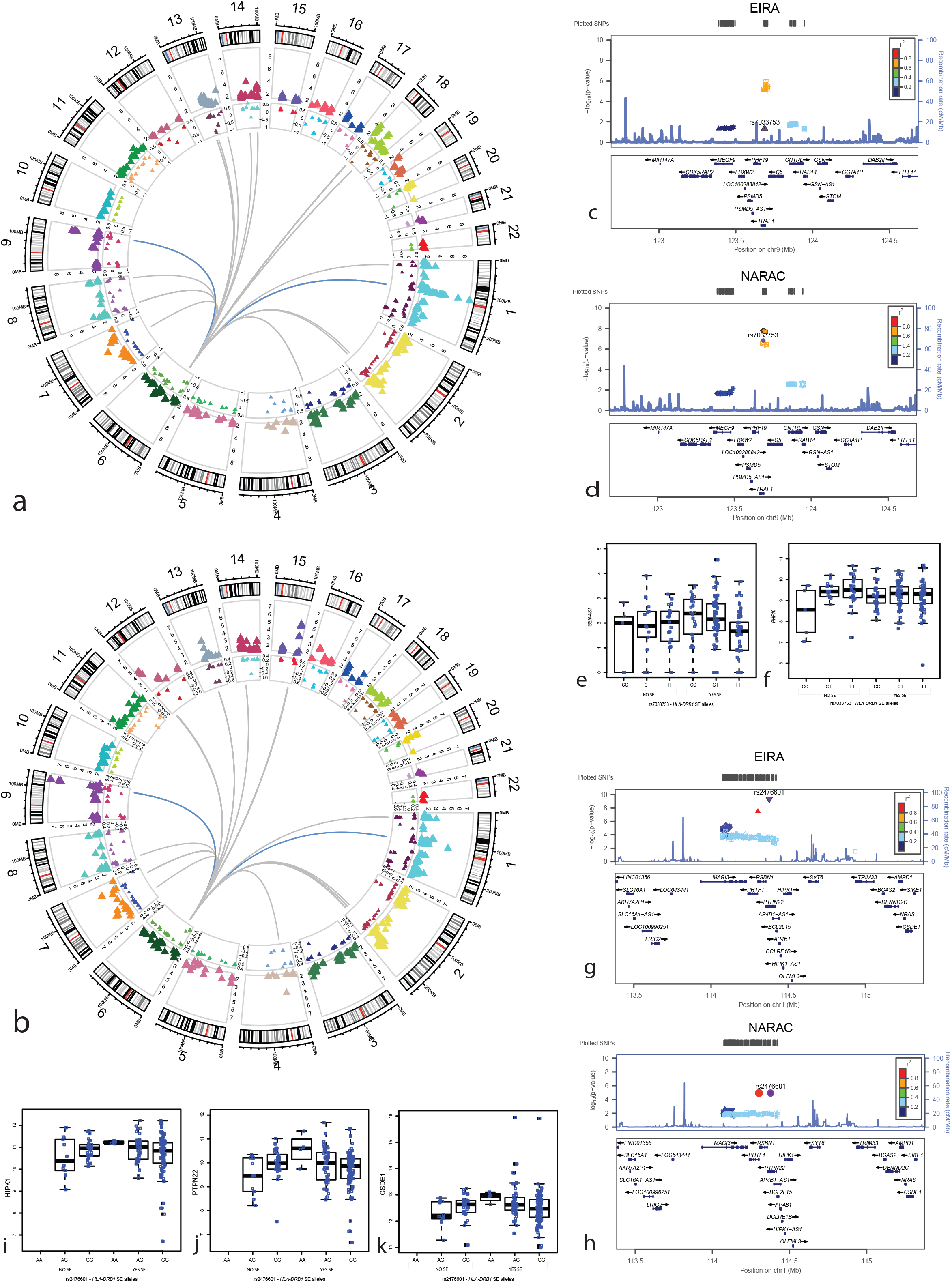
Selected SNPs from both studied cohorts with AP p-value <0.05 and same direction of AP for the additive interaction test with the *HLA-DRB1* SE alleles. The circos plots for **(a)** the EIRA study and **(b)** the NARAC study represent with triangles each of the 1,492 selected SNPs in additive interaction with the *HLA-DRB1* SE alleles. The outermost track of the circos plots is the cytoband for 22 human chromosomes. The y-axis of the second track is the negative logarithm of the AP p-values due to additive interaction with the *HLA-DRB1* SE alleles. In the third track, the y-axis corresponds to the AP value. The internal connector lines highlight the interactions that exhibited an AP p-value<1e-03. **(c-d)** Representation of locus at chromosome 9q33 centered on rs7033753 SNP for **(c)** the EIRA study and **(d)** the NARAC study. The rs7033753 is in LD (r^2^>0.6) with several other variants in interaction with the *HLA-DRB1* SE alleles in both the studies. **(e-f)** The genotype of rs7033753 variant is significantly correlated with the expression of **(e)** *GsN-AS1* and **(f)** *PHF19* genes in PBMCs from the ACPA-positive RA patients when stratification by the *HLA-DRB1* SE allelic status is considered (SE-eQTL FDR q-value=0.04 for both SNP-gene pairs). **(g-h)** Representation of locus at chromosome 1p13 centered on rs2476601 SNP for **(c)** the EIRA study and **(d)** the NARAC study. **(i-k)** The rs2476601 genotype in stratification by the *HLA-DRB1* SE allelic status significantly correlates with **(i)** *HIPK1*, **(j)** *PTPN22*, and **(k)** *CSDE1* genes expression in PBMCs from the ACPA– positive RA patients (SE-eQTL FDR q-value=0.04). Panels (c), (d), (g) and (h) were done using LocusZoom(v0.4.8) (<u>http://locuszoom.org/genform.php?type=yourdata</u>)(53, 54). Abbreviations: EIRA – epidemiological investigation of rheumatoid arthritis, NARAC – North American rheumatoid arthritis consortium, AP – attributable proportion due to interaction, LD – linkage disequilibrium, PBMCs – peripheral blood mononuclear cells, ACPA-positive RA – anti-citrullinated protein antibodies positive rheumatoid arthritis, SE-eQTL – expression quantitative trait loci in shared epitope context, FDR – false discovery rate. *GSN-AS1, PHF19, HIPK1, PTPN22*, and *CSDE1* are abbreviations for the genes.

The loci 9q33 (Fig. 4c and Fig. 4d) and 1p13 (Fig. 4g and Fig. 4h) contain the SNPs in interaction with the *HLA-DRB1* SE alleles that exhibited the lowest AP p-values (Supplementary Material Table S3). Several SNPs in interaction with the *HLA-DRB1* SE alleles from the 9q33 locus are in moderate LD (r^2^≥0.6≤0.8) among them (Fig. 4c and Fig. 4d, Supplementary Material Table S3). For instance, the rs3761847 SNP is one of the top replicated variants (EIRA: AP=0.38, 95%CI=0.22–0.55, AP p-value=6.9e-6, FDR q-value=0.04; NARAC: AP=0.43, 95%CI=0.29–0.59, AP p-value=1.6e-8, FDR q-value=2.5e-4, Supplementary Material Table S3) which has previously been associated with RA (3, 5, 17, 21, 22), and it is in moderated LD (r^2^=0.73) with the rs7033753 SNP, which is an SE-eQTL for the *GSN-AS1* (GSN antisense RNA 1) and *PHF19* (PHD finger protein 19) genes (Fig. 4f to Fig. 4g, Supplementary Material Table S5). On the other hand, the top SNP in interaction with the *HLA-DRB1* SE alleles is the non-synonymous variant rs2476601 in the *PTPN22* gene located in the 1p13 locus (Fig. 4g and Fig. 4h). Although the interaction between rs2476601 SNP and the *HLA-DRB1* SE alleles in ACPA-positive RA has been reported previously (7), we observed in our SE-eQTLs analysis that the rs2476601 SNP is an SE-eQTL for *HIPK1* (homeodomain interacting protein kinase 1), *PTPN22* (protein tyrosine phosphatase, non-receptor type 22), and *CSDE1* (cold shock domain containing E1) genes (Fig. 4i to Fig. 4k, Supplementary Material Table S5). Additionally, there is evidence from capture Hi-C technology that the rs2476601 physically interacts with the *HIPK1* and *CSDE1* genes in foetal thymus cells, monocytes, CD4 naïve T cells, CD8 naïve T cells, neutrophils and B cells (https://www.chicp.org)(23–25). This supports our finding of rs2476601 SNP as SE-eQTLs and suggest that the detected additive interaction with *HLA-DRB1* SE alleles in ACPA-positive RA likely reflect functional implication.

We applied gene ontology (GO) analyses of these 1,492 selected SNPs and 56 terms where highlighted as significant after FDR correction (Supplementary Material Table S6). The list of significant GO terms, when ranked by the FDR hypergeometric q-value, is enriched by pathways related to regulation of secretion (5 terms), signaling (11 terms), cell differentiation and development (11 terms), immune cells related (3 terms) and bone disease related (5 terms), which are relevant to ACPA-positive RA.

Together, these results suggest the plausibility of the high impact of 1,492 selected SNPs for the pathogenesis of ACPA-positive RA through interaction with the *HLA-DRB1* SE alleles. Nevertheless, when multiple testing correction by FDR was applied 15 SNPs remain significant (from the 1p13 and 9q33 loci; AP FRD q-value < 0.05) in both EIRA and NARAC (Supplementary Material Table S3). Thus, these results require additional replication in independent cohorts of ACPA-positive RA patients and controls.

Finally, we observed that the step-by-step removal of the risk alleles of the two-top replicated SNPs in interaction with the *HLA-DRB1* SE alleles (rs2476601 at 1p13 and rs10739581 at 9q33, AP FDR q-value<0.05), decreases the effect size of SE alleles for ACPA positive RA in the studied cohorts (Fig 5). This observation also suggests that the association between the *HLA-DRB1* SE alleles and the risk of ACPA-positive RA is at least partly influenced by multiple interactions with non-HLA genetic variants.

**Fig 5.**
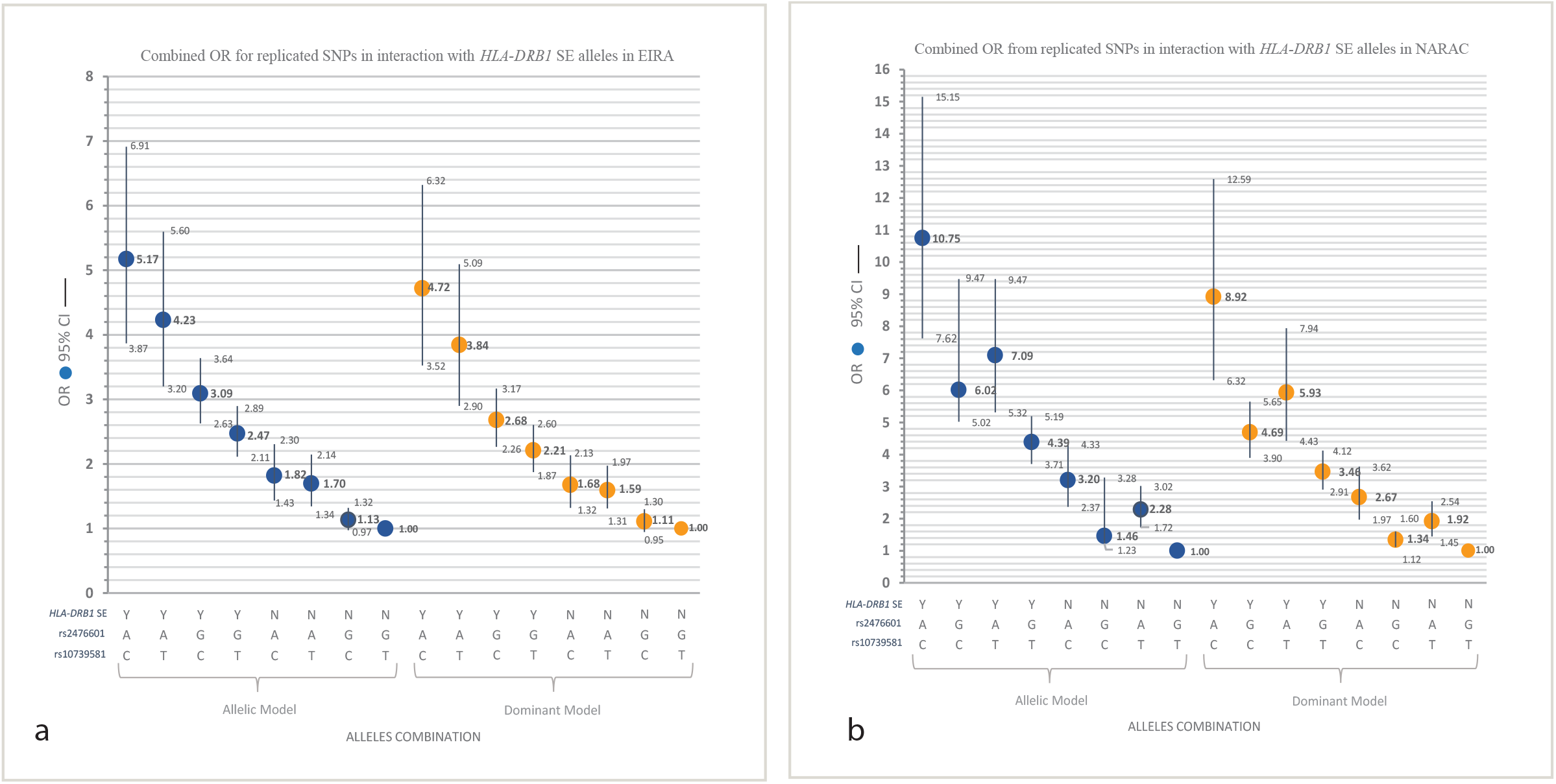
Three-factor’s OR calculation: the *HLA-DRB1* SE alleles and two of the replicated SNPs in significant interaction. On the x-axis – the combinations of presence or absence of risk alleles for allelic or dominant models. On the y-axis – the combined ORs with 95% CI of *HLA-DRB1* SE alleles (presence – Y, or absence – N), the rs2476601(G>A, in the 1p13 locus) SNP and the rs10739581(T>C, in the 9q33 locus). The YAC allelic combination is a risk factor to develop ACPA-positive RA in the study populations. Panel **(a)** shows data from EIRA study, where both rs2476601 and rs10739581 are in significant interaction with the *HLA-DRB1* SE alleles after FDR correction (AP=0.45 95%CI=0.31–0.60, p-value=4.3e-10, FDR q-value=5.2e-5 and AP=0.40 95%CI=0.24–0.57, p-value=1.4e-6, FDR q-value=0.04, respectively). Similarly, panel **(b)** shows data from NARAC study for the combined OR of *HLA-DRB1* SE alleles, rs2476601 and rs10739581 variants. The rs2476601 SNP at 1p13 locus and rs10739581 at 9q33 locus are in significant interaction with *HLA-DRB1* SE alleles after FDR correction in the NARAC study (AP=0.41 95%CI=0.23–0.6, p-value=1.1e-5, FDR q-value=0.04 and AP=0.43 95%CI=0.28–0.6, p-value=2.1e-8, FDR q-value=2.5e-4, respectively). Abbreviations: OR – odds ratio, CI – confidence intervals, ACPA-positive RA – anticitrullinated protein antibodies positive rheumatoid arthritis, EIRA – epidemiological investigation of rheumatoid arthritis, NARAC – North American rheumatoid arthritis consortium, AP – attributable proportion due to interaction, FDR – false discovery rate.

## Discussion

Our study of two independent ACPA-positive RA cohorts demonstrates that the *HLA-DRB1* SE alleles are involved in multiple interactions with disease-associated SNPs in comparison to nonassociated SNPs. We show evidence of gradual decrease of the effect size of the *HLA-DRB1* SE alleles in the risk of ACPA-positive RA after adjusting for top SNPs in interaction (Figs. 3 and 5). Based on these findings, we would like to propose the *sovereignty hypothesis*, which suggests that the *HLA-DRB1* SE alleles act as a genetic hub of simultaneous multiple interactions with the non-HLA genetic variants that by themselves have a modest effect size in RA (OR<2), and in turn, cumulatively contribute to the high effect size of the *HLA-DRB1* SE alleles in development of the ACPA-positive RA. Our hypothesis deals with a missing link that integrates the *HLA* alleles with other genetic variations across the human genome in providing knowledge about the risk of developing this common autoimmune disease.

Low statistical power and inevitable high number of type I and type II errors hamper the genome-wide analysis of the gene-gene interactions in existing RA cohorts. Therefore, we chose to address the distribution of probabilities associated with the interaction statistics, AP (attributable proportion due to interaction), using a comparison between empirically observed ACPA-positive RA risk (nominal association p-values <0.05) and non-risk SNPs (nominal association p-values ≥0.05). With this relatively liberal threshold, we can expect that the first group will be enriched with true ACPA-positive RA associated variations, while the second group will be enriched with true non-associated variations. With this setup, our results clearly indicate that there is a strong difference in the distribution of p-values of interaction (AP p-values) between both groups of predefined SNPs, and this difference is mainly due to an enrichment of small p-values of interaction between the *HLA-DRB1* SE alleles and the ACPA-positive RA associated polymorphisms. These observations are in line with our *sovereignty hypothesis*, which also has foundations in the sufficient-component cause model (14). This model suggests that diverse components are part of a sufficient cause for a disease in a given affected individual, where each sufficient cause can include one or more component causes and form a minimal set of conditions that yield disease (26). Our study demonstrated that the *HLA-DRB1* SE alleles are a relevant component (but non-sufficient by itself) in the cause of ACPA-positive RA by interacting with multiple non-essential genetic risk factors.

Interestingly, a study showed that interacting loci were part of radial epistatic networks, where the hub loci interacted with multiple quantitative trait loci (QTLs); the hub locus acts as a genetic capacitor that modifies the effect of the radial loci in the network (27). If we extrapolate Forsberg *et al.’s* (27) observations with our results, we could assume that in ACPA-positive RA the principal QTL hub is represented by the *HLA-DRB1* SE alleles. This hub concentrates a complex interaction network of multiple non-HLA QTLs, where the effects could be modified mutually.

Interactions between the variants of the *HLA* region in ACPA-positive RA could be expected and this together with the strong LD in this locus prompted us to exclude the extended *MHC* region from our study, for the sake of simplicity. A previous study has explored interactions among the different *HLA* alleles (8), nevertheless a more extended investigation of these intricate interactions is required.

The present finding of multiple polymorphisms interacting with the *HLA-DRB1* SE alleles and their mutual effect size influence in the risk for disease, suggest that many mechanisms will affect the impact of *HLA-DRB1* SE alleles in the context of ACPA-positive RA. Although the present analysis was mainly performed with statistical methods, some preliminary functional mechanisms can be extrapolated from our data. Indeed, the statistical approach in our study resulted in a list of 1,492 SNPs as good candidates that interact with the *HLA-DRB1* SE alleles in the risk of developing of ACPA-positive RA. From them, 13.5% are suggested SE-eQTLs in ACPA-positive RA individuals, indicating that the additive interactions detected may be a reflection of biological processes.

For instance, ACPA-positive RA patients who carry both the risk allele of the top interacting variant, the rs2476601 SNP, and the *HLA-DRB1* SE alleles, seem to have a higher expression of *PTPN22, HIPK1* and *CSDE1* genes in PBMCs (Figs. 4i to 4k). Intriguingly, the rs2476601 SNP physically interacts with the *HIPK1* and *CSDE1* genes in certain type of immune cells, including CD4+ T cells, that in turn are known to be relevant in the pathogenesis of RA (28). Moreover, the T-box transcription factor Eomesodermin *(EOMES)* is part of 11 highlighted GO terms in our analysis (Supplementary Material Table S6). *EOMES* was annotated due to three SNPs in interaction with *HLA-DRB1* SE alleles (rs1506691, rs6804917 and rs12630663; Supplementary Material Tables S3 and S6), which interestingly also physically interact with *EOMES* in CD4+ and CD8+ T cells (https://www.chicp.org)(23, 25). EOMES is a transcription factor important for memory T cell formation and cytotoxic T cell differentiation (29). On the other hand, a study has demonstrated that MHC genotype, and *HLA-DRB1* in particular, has a key role in shaping the T cell receptor repertoire (30), evidence that goes in line with our suggestion that the observed statistical interactions are a reflection of functional implications. Nevertheless, additional replication and interpretation of these interactions (pointed by the 1,492 selected SNPs and the highlighted GO terms centered in the *HLA-DRB1* locus) in relation to biological processes will be the next step to further increase the etiological understanding of ACPA-positive RA.

The four most statistically significant SNP-gene pairs from the SE-eQTL analysis are new candidates in the genetic component of RA: rs10404242-TLE6 (transducing like enhancer of split 6), rs576363 8-ZNRF3-AS1 (ZNRF3 antisense RNA 1), rs28513183-HSD11B1 (hydroxysteroid 11-beta dehydrogenase 1), and rs1781279-MTPAP (mitochondrial poly(A) polymerase). Particularly, the *HSD11B1* gene encodes the 11ß-HSD1 enzyme involved in the biosynthesis of steroid hormones, related to an increase in the intracellular glucocorticoids levels. A knockdown *HSD11B1* mice model presents an increased acute inflammation after induction of experimental arthritis (31). Notably, all these SNPs show eQTL effects only in a context of the *HLA-DRB1* SE alleles, and further implication in ACPA-positive RA should be elucidated.

In conclusion, we used a new approach for the investigation of interactions at the genome-wide level in ACPA-positive RA, which led us to detect a significant enrichment of interactions between the *HLA-DRB1* SE alleles and associated with the disease SNPs in comparison to all other, non-associated SNPs. There is a visible reduction of the size of the effect of *HLA-DRB1* SE alleles on ACPA-positive RA risk when the risk alleles of the top interacting SNPs are discounted in a combined OR calculation (Fig 5). Our approach is potentially applicable to other autoimmune diseases or complex traits, where a single or a limited number of strong risk factors are observed. This approach could be used as a tool to explore the next level of complexity of multifactorial diseases, eventually allowing the detection of interconnected genetic variants in the risk for a phenotype, which could in turn contribute to a better understanding of disease mechanisms.

## Materials and Methods

### Studied populations

This project was based on GWAS data from two independent case control studies of RA, EIRA (5, 12, 15–18) and NARAC (5, 16, 19, 20). Briefly, the EIRA study recruited incident RA cases and healthy individuals selected from a national register that matched the cases by gender, age, and residence area (17, 18). Unrelated RA cases from multicase families from the United States were included in the NARAC study and matched with unrelated controls recruited from the New York Cancer Project (17, 19). In both studies, the RA patients were diagnosed based on the American College of Rheumatology (ACR) criteria from 1987 (32). Ethical approval was guaranteed for each study from the respective ethical committees and are in accordance with the Declaration of Helsinki. A total of 4,291 individuals were included in this study, with 1,151 ACPA-positive RA cases and 1,079 healthy controls from EIRA and 867 ACPA-positive RA cases and 1,194 healthy individuals from NARAC (Table 1).

### HLA genotyping

*HLA* typing in the EIRA study was made by sequence-specific primer polymerase chain reaction assay (SSP-PCR) (DR low-resolution kit; Olerup SSP, Saltsjöbaden, Sweden), and the PCR products were loaded into 2% agarose gels for electrophoresis. An interpretation table was used to determine the specific genotype according to the manufacturer’s instructions (33). In the NARAC study, the *HLA* typing was also performed by SSP-PCR based methods as described elsewhere (34).

*HLA-DRB1* SE alleles included *01 (except *0103), *04 (using high resolution data for *0404, *0405 and *0408 when possible), and *1001. A variable for the *HLA-DRB1* SE alleles was coded as NN, NY, and YY genotype like, where N and Y stand for “no” or “yes” based on the presence of the *HLA-DRB1* SE alleles.

### GWAS data, data filtering, and SNP grouping

As described previously (5), the genotyping platforms used were HumanHap300 BeadChip and HumanHap550 BeadChip from Illumina^®^ for EIRA and NARAC, respectively. The data were filtered for minor allele frequency (MAF) <1%, missing rate higher or equal to 5%, and p-values <0.001 for Hardy-Weinberg equilibrium (HWE). A principal component analysis (PCA) was performed using the EIGENSOFT (v6.1.1) (<u>https://www.hsph.harvard.edu/alkes-price/software/</u>)(35) software to model the population stratification between the cases and controls after removing the extended *MHC* region and pruning the GWAS data sets (from nonimputed SNPs) based on the linkage disequilibrium (LD), excluding a SNP from a pair when their r^2^ was higher than 0.5. The 1000 Genomes Phase I (α) Europeans was used as a reference panel for imputation in IMPUTE2 (v2.3.0) (<u>https://mathgen.stats.ox.ac.uk/impute/impute_v2.html#home</u>)(36) for EIRA and minimac (release stamp 2011–10–27) (<u>http://genome.sph.umich.edu/wiki/Minimac</u>)(37) for NARAC. Duplicated SNPs and SNPs with a low imputation score (Rsq <0.5) were removed; thereafter, the same filters of MAF, missing rate, and HWE, described above were applied again for both cohorts. The sex chromosomes were not included in the present study.

We removed the extended *MHC* region (chr6:27339429 to chr6:34586722, hg19) from on our analyses, to exclude the influence of high LD and independent signals of association (38). A logistic regression model implemented in plink (v1.07) (<u>http://pngu.mgh.harvard.edu/~purcell/plink/</u>)(39) was used to estimate the association between each of the SNPs in GWAS and risk of ACPA-positive RA in EIRA and NARAC. Based on the nominal p-values of association, the SNPs were grouped into risk (p-values <0.05) or nonrisk SNPs (p-values ≥0.05). The number of SNPs and percentages are shown in Fig 1b and Table 2. Five percent of the EIRA imputed SNPs showed a nominal p-value of association less than 0.05, while 8.7% was observed in the NARAC. There was an overlap of 19,769 SNPs between the two studies.

### Interaction Analysis

After applying the filters mentioned above, we tested for additive interaction between the *HLA-DRB1* SE and each SNP from the EIRA and NARAC GWAS. The null hypothesis of the additive model assumes that there is additivity between the different sufficient causes for a phenotype, while the alternative hypothesis is assumed when departure from additivity is observed. The departure from additivity is estimated by the attributable proportion (AP) due to interaction using OR as the risk estimates(40) with the following equation:

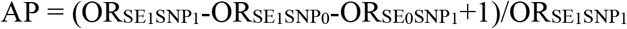

Where 1 and 0 refer to presence or absence of the risk factor/allele respectively, the ORs are calculated using SE0SNP0 as a reference group. A cut-off of five for each of the cell frequencies was applied in the interaction analysis. The gender and the first ten principal components from PCA were included as covariates in the model. AP value, its respective p-value and confidence interval (95%CI) were assessed using logistic regression by means of the program GEISA(v0.1.12) (<u>https://github.com/menzzana/geisa</u>)(11, 41). An update JAVA coded software of the previously published GEIRA algorithm (42). The numbers and the percentage of SNPs analyzed for each studied cohort are presented in Table 2.

### Comparison of the distribution of AP p-values between the risk and non-risk groups of SNPs and quality control approaches

The distribution of AP p-values observed in the interaction analysis from the ACPA-positive RA risk SNPs was compared with the distribution of AP p-values observed in the interaction analysis from the non-risk SNPs using the Kolmogorov-Smirnov (KS) test, implemented in the *stats* package of R software(v3.3.2) (https://www.r-project.org/)(43). The KS test statistic quantifies the maximum distance (D) between the two empirical cumulative distribution functions (ECDF) of the AP p-values from the risk and non-risk SNPs groups. The alternative hypothesis for the KS test was that the ECDF of the AP p-values from the risk SNPs is higher than the one for the non-risk SNPs influenced by an enrichment of small AP p-values due to the interaction between the *HLA-DRB1* SE alleles and the risk group of SNPs. The p-value obtained from this KS test was lower than the machine precision, represented as <2.2e-16 or zero when the absolute p-value was asked. The <2.2e-16 value corresponds to the default *double.eps* component of the numerical characteristics of R machine. Thus, the threshold for the permutations was set to 2.2e-16. We permuted the category for the SNPs, of risk and nonrisk ten thousand times, applying the KS test each time to identify the proportion of p-values from the KS tests that are less than the set threshold (<2.2e-16). The percentage of the KS test results with p-values less than 2.2e-16 was 0, the maximum D value observed was 6.1e-03 in both analyzed cohorts. Likewise, we permuted the *HLA-DRB1* SE variable using non-imputed GWAS data and a smaller number of permutations (n=1000), due to the computational limitations to calculate the interaction for each randomized SE variable against all SNPs in the GWAS. We applied the KS test to detect differences in the AP p-values’ distribution for risk (nominal p-values of association <0.05) versus non-risk (nominal p-values of association ≥0.05) SNPs, each time the SE variable was randomized. The maximum D values observed were 0.05 and 0.03 for EIRA and NARAC, respectively. The percentage of the KS test p-values from permutations less than 2.2e-16 were 0.1 for both EIRA and NARAC. Both types of permutations showed that less than 5% of the KS test will exhibit a p-value under 2.2e-16, strongly indicating that differences in the AP p-values distribution detected by the KS test from the original data are unlikely to be by chance.

In order to verify that the observed results were not due to statistical artifacts, we employed several approaches. First, we performed two types of permutations, for SE alleles and for the groups of SNPs, described in detail above. Second, we removed the SNPs from the *PTPN22* locus (chr1:113679091 to chr1:114679090, GRCh37/hg19) and applied the same workflow, from step one to nine of Fig 1b, to determine whether this locus significantly influences the observed enrichment due to the known gene-gene interaction between the rs2476601variant (or SNPs in LD with this variant) and the *HLA-DRB1* SE alleles (7). Third, we used the above mentioned workflow, from step one to nine of Fig 1b, replacing the SE variable with the rs4507692 SNP as a negative control, since the rs4507692 SNP is not associated with RA but has the same MAF as the *HLA-DRB1* SE alleles (Table 1). Fourth, non-imputed GWAS data was used in the same methodological workflow, from step one to nine of Fig 1b, as well as removing data from the entire chromosome 6, to address possible inflation in the results due to a high LD with the *HLA-DRB1* SE alleles.

### SNPs in interaction with SE between EIRA and NARAC

We selected those variants with AP p-values < 0.05 and same AP direction from both the EIRA and NARAC studies to evaluate their distribution across the genome and their possible implication in expression of the neighboring genes in the *HLA-DRB1* SE alleles context. We also applied FDR correction to the AP p-value of these 1,492 SNPs and considered significant q-values less than 0.05 (Supplementary Material Table S3).

### Expression Quantitative Trait Loci (eQTL) in the context of the HLA-DRB1 SE alleles

We evaluated whether the selected SNPs in interaction with the *HLA-DRB1* SE alleles were eQTLs in the *HLA-DRB1* SE alleles context for genes ±1Mbp around them (GRCh37/hg19 assembly was used) using data from the PBMCs found in the COMBINE study (44). Briefly, the PBMCs from the RA patients were sampled at the Rheumatology Unit, Karolinska Institute, Stockholm-Sweden. RNA was purified from the PBMC samples and sequenced using Illumina HiSeq 2000 with TruSeq RNA sample preparation. DNA was genotyped using Illumina OmniExpress arrays (12v1) (44).

The eQTLs in the context of the *HLA-DRB1* SE alleles (SE-eQTLs) were analyzed in 97 ACPA-positive RA patients (69% females) that were undergoing a change or start of a new treatment regimen. In the analysis, the following formula was applied:

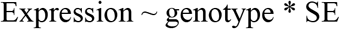

Where expression was the normalized gene expression values of a proximal gene, processed according to the edgeR package (v3.8.6) (https://bioconductor.org/packages/release/bioc/html/edgeR.html)(45, 46) i.e., the log2 transformed and TMM-normalized (trimmed mean of M-values normalization method). The genotype was the numerically encoded genotype of the SNP (AA=0, AB=1, BB=2), and SE was the *HLA-DRB1* shared epitope alleles status as either true or false. Sex and treatment cohort were used as covariates and subject-ID as repeated measure, which was assumed as random intercept for each subject in the mixed-linear model (Data available on request). The calculation was performed using the mixed-linear model function in the nlme 3.1 package from R/Bioconductor (v3.3.2) (https://www.bioconductor.org/)(47). Low expressed genes (TMM-normalized < 1.4) were filtered out and 5% FDR(48) was considered.

### Gene ontology analysis

In order to have a global view of the plausible biological pathways pointed by these SNPs in interaction with the *HLA-DRB1* SE alleles, gene ontology (GO) was assessed with GREAT (v3.0.0) (<u>http://bejerano.stanford.edu/great/public/html/index.php</u>)(49) to the list of 1,492 SNPs. Those SNPs have AP p-values <0.05 and same AP direction in both the EIRA and NARAC results (Supplementary Material Tables S6 and S7).

## Acknowledgments

We would like to thank all the patients and control individuals, involved in the EIRA, NARAC and COMBINE studies. Thanks to Magdalena Lindén, Tojo James, and Ingrid Kockum, whom provided us an updated version of *GEISA* and good discussions about this tool. Thanks to Soumya Raychouhundry and Peter Gregersen, that provided us with continuous help and supported us with data from NARAC. Thanks to Gilad Silberberg, who gave input in redaction and discussion of the results. We would also like to thank The National Genomics Infrastructure (NGI) in Sweden for providing computational resource for our study and Meena Strömqvist for English language editing.

## Supplementary Material

**Fig S1. Comparison of the distribution of p-values for attributable proportion in the EIRA and NARAC studies for interaction tests between the SNP rs4507692 and RA risk or nonrisk SNPs.** The rs4507692 is a variant that has the same MAF as the *HLA-DRB1* SE alleles but is not associated with ACPA-positive RA. **(a)** Density plot of the AP p-values for the interaction between rs4507692 and the ACPA-positive RA risk group of SNPs (raw p-value of association <0.05) or **(b)** non-risk group of SNPs (raw p-value of association ?0.05) in the EIRA study. **(c)** The respective ECDF plot of the AP p-values distribution of risk (red line) or non-risk (blue line) SNPs in interaction with the rs4507692 (KS test, D=0.018, p-value=1.4e-43, (Table 1 and Supplementary Material Tables S1 and S2). We tested for differences in the AP p-values distribution on the fraction that could be considered as significant interactions with rs4507692 (AP p-value <0.05). **(d)** Density plot for the AP p-values from the interaction tests between the risk SNPs and rs4507692 or, **(e)** between the non-risk SNPs and rs4507962 SNP in the EIRA study. **(f)** ECDF of the fraction of the AP p-values distribution corresponding to <0.05 in the EIRA study (KS test, D=0.009 and p-value= 0.50). Similar results were observed from the NARAC study, an independent replication cohort: **(g)** Density plot of the AP p-values for the interaction between the rs4507692 and the risk group of SNPs or **(h)** the non-risk group of SNPs. **(i)** The respective, ECDF plot from the NARAC study (KS test, D=0.001, p-value= 0.458, Supplementary Material Table S2). **(j)** Density plot of the fraction of the AP p-values of less than 0.05 from the interactions between the rs4507692 and the risk SNPs or (k) the nonrisk SNPs. **(l)** The ECDF plot from this fraction of the AP p-values distribution (KS test, D= 0.027 and p-value=1.29e-07) is not significant since it is higher than the significant threshold set for the KS-test of 2.2e-16.

Abbreviations: EIRA – epidemiological investigation of rheumatoid arthritis, NARAC – North American rheumatoid arthritis consortium, ACPA-positive RA – anti-citrullinated protein antibodies positive rheumatoid arthritis, AP – attributable proportion due to interaction, ECDF – Empirical cumulative distribution function, KS test – Kolmogorov – Smirnov test.

**S2 Figure. ECDF for the KS test between the AP p-values of the risk SNPs test for interaction with the *HLA-DRB1* SE alleles (upper line in light red) or with the rs4507692 variant (bottom line in dark red). (a)** in EIRA (KS test, D=0.352, p-value< 2.2e-16) and **(b)** in NARAC (KS test, D=0.258, p-value<2.2e-16).

Abbreviations: ECDF – Empirical cumulative distribution function, KS test – Kolmogorov – Smirnov test, AP – attributable proportion due to interaction.

**S3 Figure. The four most significant SE-eQTLs**. SNPs in interaction with the *HLA-DRB1* SE alleles from the EIRA and NARAC studies were selected (AP p-value <0.05 and same direction of AP in both studies) and evaluated as cis-eQTL in the presence or absence of the *HLA-DRB1* SE alleles (SE-eQTL) in PBMCs from the ACPA-positive RA patients (COMBINE study(44)). We observed that 201 SNPs in interaction with the *HLA-DRB1* SE alleles are eQTLs when the SE allelic status is considered (FDR q-value<0.05). The top four SNP-gene pairs are represented in the plots: **(a)** rs10404242-*TLE6*, SNP-gene pair (SE-eQTL p-value=6.7e-4, FDR q-value=0.04). The rs10404242 variant is in interaction with the *HLA-DRB1* SE alleles in the EIRA study (AP= 0.25, 95%CI=0.09–0.42, AP p-value=0.002) and in the NARAC study (AP=0.23 95%CI=0.04–0.43 AP p-value=0.02). **(b)** *rs5763638-ZNRF3-AS1*, SNP-gene pair (SE-eQTL p-value=1.9e-3, FDR q-value=0.04). The rs5763638 variant is in interaction with the *HLA-DRB1* SE alleles in the EIRA study (AP=0.19, 95%CI=0.013–0.37, AP p-value=0.03) and in the NARAC study (AP=0.22, 95%CI=0.02–0.44, AP p-value=0.03). **(c)** rs28513183-*HSD11B1*, SNP-gene pair (SE-eQTL p-value=2e-3, FDR q-value=0.04). The rs28513183 variant is in interaction with the *HLA-DRB1* SE alleles in the EIRA study (AP=0.22, 95CI=0.02–0.44, AP p-value=0.03) and in the NARAC study (AP=0.27, 95CI=0.07–0.49, AP p-value=9.4e-3). **(d)** rs1781279-MTPA, SNP-gene pair (SE-eQTL p-value=2.9e-3, FDR q-value=0.04). The rs1781279 variant is in interaction with the *HLA-DRB1* SE alleles in the EIRA study (AP=0.19, 95%CI=0.02–0.37, AP p-value=0.03) and in the NARAC study (AP=0.24, 95%CI=0.05–0.44, AP p-value=0.01).

Abbreviations: SE-eQTL – expression quantitative trait loci in shared epitope context, EIRA – epidemiological investigation of rheumatoid arthritis, NARAC – North American rheumatoid arthritis consortium, AP – attributable proportion due to interaction, PBMCs – peripheral blood mononuclear cells, ACPA-positive RA – anti-citrullinated protein antibodies positive rheumatoid arthritis, FDR – false discovery rate. *TLE6, ZNRF3-AS1, HSD11B1* and *MTPA* are abbreviations for the genes.

**Table S1.** The Kolmogorov-Smirnov (KS) test for AP p-values distributions of the interaction analysis with the rs4507692 SNP ^a, b^ in EIRA and NARAC imputed data.

**Table S2.** The Kolmogorov-Smirnov (KS) test for AP p—values distributions of the interaction analysis in EIRA and NARAC GWAS (non-imputed data).

**Table S3.** Selected SNPs in interaction with the *HLA-DRB1* SE alleles from EIRA and NARAC. The SNPs were selected whether they exhibited AP p-values < 0.05 and the same direction of AP in both studies. ^a^ The interaction tests were done using the risk allele from the tested SNPs.^b^ 1 refers to the risk factor or alleles. Complementary 0 refers to the opposite no-risk factor or allele.

**Table S4**. Distribution across the human genome of the selected SNPs in interaction with the *HLA-DRB1* SE alleles.

**Table S5**. SE-eQTLs observed in ACPA-positive RA patients. Only selected SNPs in interaction with the *HLA-DRB1* SE alleles were tested.

**Table S6**. Gene ontology (GO) terms obtained using 1,492 selected SNPs in interaction with the *HLA-DRB1* SE alleles.

**Table S7**. Setting used, input, output data, and results from the gene ontology (GO) analyses.

